# Molecular and evolutionary determinants for protein interaction within a class II aldolase/Adducin domain

**DOI:** 10.1101/2024.12.18.629236

**Authors:** Marina E. Seheon, Amalia S. Parra, Christopher A. Johnston

**Author notes:** these authors contributed equally to this work (Marina E. Seheon, Amalia S. Parra).

## Abstract

The appearance of modular protein interaction domains represents a crucial step in the evolution of multicellularity. For example, the class II aldolase domain (ALDO^DOM^) found within the Adducin gene family shares sequence and structural homology to a glycolytic aldolase enzyme found in many evolutionarily ancient phyla. ALDO^DOM^ is best known for direct binding to actin filaments through a tetrameric assembly. Molecular details for additional ALDO^DOM^ interactions have not been resolved, nor have the sequence changes underlying the dramatic functional switch in the aldolase protein fold. Here we explore the molecular basis for the interaction between ALDO^DOM^ of Hts (*Drosophila* Adducin) and the mitotic spindle regulator, Mud. Our results suggest a distinct mode of interaction, as conserved actin-contacting residues on the tetramer surface were found dispensable for Mud binding. Instead, we identify a critical role for the ALDO^DOM^ C-terminal helix (CThelix), along with residues from the adjacent protomer that occur at a tetrameric interface conserved among domains and a subgroup of aldolase enzymes (ALDO^ENZs^). Truncation of the CThelix from bacterial ALDO^ENZ^, or chimeric fusion with that from Hts, confers ALDO^DOM^-like Mud binding. Sequence database analyses suggest ALDO^DOM^ function may have arisen in the primitive metazoan phylum, *Placozoa*, which contain both an aldolase enzyme and domain capable of Mud binding. Finally, we identify a single, conserved arginine-to-glycine change that also permits Mud binding within the bacterial ALDO^ENZ^. Our work provides molecular and evolutionary insights into the function of a conserved protein-binding domain within multicellular organisms.

## INTRODUCTION

Acquisition of new protein functions has been an important feature in the evolution of multidomain proteins typical of multicellular organism proteomes. In many cases, this neofunctionalization is thought to occur following gene duplication events, with diverging sequences leading to changes in globular structure, conformational dynamics, protein-protein interaction surfaces, and domain combinations that ultimately adopt new functions often at the expense of the previous known function (1, 2). Another driver of protein evolution involves the co-option of existing genes to perform novel functions following diverging sequence changes (3). Together, these events lead to gene evolution that generate complex functions in multicellular life such as cell signaling, cell adhesion, and differentiation. However, very little is known about the underlying molecular mechanisms that establish newly evolved protein functions.

One recent example addressing this issue is the evolution of guanylate kinase (GK), an ancient enzyme involved in nucleotide metabolism, into a protein binding domain conserved across metazoan phyla (4–7). While studies on some enzymes have described relatively minor functional innovation such as substrate preference (8), studies on GK evolution describe a major shift in function, that from a catalytic enzyme to a protein interaction platform. Notably, this functional change occurred while preserving the substrate-binding site in the GK enzyme, which was co-opted as the primary site of protein interaction in GK domains (5). Sequence divergence led to restrictions in GK allostery necessary for productive protein-protein interactions (5, 7).

A strikingly similar functional switch may have occurred in the class II aldolase protein fold. Current theories suggest the glycolytic aldolase enzyme (ALDO^ENZ^) lost catalytic activity as it was integrated into the Adducin family of cytoskeletal proteins widely conserved across multicellular life. Adducin plays a critical role in organizing cortical spectrin-actin structures to facilitate a wide range of biological functions (9, 10). The exact roles of the ALDO^DOM^ remain poorly defined, and a molecular basis for its shift from a standalone enzyme to a component of the larger, multidomain Adducin protein has not been rigorously addressed. Recently, cryo-EM structural analysis of the erythrocyte spectrin-actin complex revealed that the Adducin ALDO^DOM^ forms a tetramer that directly interacts with the barbed end of the actin filament, with additional contacts occurring via extended sequences N- and C-terminal to the core ALDO^DOM^ (11). These may not be distinguishing properties of the ALDO^DOM^, however, as several ALDO^ENZ^ structures reveal a similar tetrameric assembly that contributes to catalytic efficiency (12, 13). ALDO^ENZ^ has also been shown to bind F-actin, although the ubiquity of this interaction across aldolase subgroups and its functional significance are less clear (13, 14). Identifying other Adducin ALDO^DOM^ protein binding partners and elucidating how such interactions form remain important research pursuits.

We recently identified a direct interaction between the ALDO^DOM^ of the *Drosophila* Adducin protein, Hu li tai shao (Hts^ALDO^), and a coiled-coil domain of the mitotic spindle regulatory protein, Mushroom body defect (Mud^CC^). Functionally, Hts impacts the phase separation dynamics of Mud *in vitro* and is required for proper spindle positioning function *in vivo* (15). Our previous work, however, did not resolve a clear molecular basis for the Hts/Mud interaction. Thus, whether this complex forms in a similar manner to that seen between Adducin and actin, or uses an alternative mode of protein binding, remains unknown. Here, we combine AIphaFold 3 (AF3) structural modeling, biochemical binding assays, and comparative sequence analysis of ALDO^ENZ^ and ALDO^DOM^ proteins across diverse phyla to formulate a molecular model for protein binding within the tetrameric class II aldolase fold, using the Hts/Mud complex as a model system. In contrast to the Adducin/actin complex, mutation of conserved residues on the Hts tetramer surface did not impair Mud binding. In contrast, alteration of the Hts^ALDO^ C-terminal helix (CThelix), which AF3 modeling revealed contains an extended loop insertion compared to known bacterial enzyme structures, strongly impacted Mud binding. Mutation of conserved CThelix-contacting residues from an adjacent protomer, which form similar contacts in Adducin and ALDO^ENZ^ tetramers, also significantly reduced Mud binding. Further analysis of bacterial ALDO^ENZs^ found that truncation of their CThelix, or chimeric replacement with that from Hts, conferred strong Mud binding to these otherwise binding-deficient enzymes. A similar ALDO^ENZ^ gain-of-function was identified following a single arginine substitution to a highly conserved glycine within ALDO^DOMs^. Finally, sequence database searches across major phyla suggests that ALDO^DOM^ may have first appeared within *Placozoa*, which were also found to have retained a putative class II ALDO^ENZ^. In contrast to bacterial ALDO^ENZ^, the *Placozoan* enzyme showed robust Mud binding that was compromised in a chimeric protein with the bacterial CThelix. Overall, our data provide a molecular model for the Hts/Mud complex and implicate a crucial regulatory role for the CThelix within the class II aldolase fold in the evolution of its function as a protein binding domain.

## MATERIALS AND METHODS

### Molecular cloning

Hts and Mud sequences were amplified from a cDNA library prepared from *Drosophila* S2 cells. Additional aldolase enzyme and domain genes were amplified from plasmids constructed through complete gene synthesis by Genewiz (Azenta Life Sciences, South Plainfield, NJ, USA). All aldolase enzyme and domain sequences were cloned into the pMAL-C5x plasmid using 5’-NdeI and 3’-SalI restriction sites. The Mud^CC^ domain was cloned into pBH with an N-terminal TEV-cleavable hexahistidine tag using 5’-BamHI and 3’-XhoI restriction sites.

### Protein purification

All proteins were expressed in BL21(DE3) *E. coli* under induction of isopropyl β-D-1-thiogalactopyranoside (IPTG) and grown in standard Luria–Bertani broth supplemented with 100 μg/ml ampicillin. Transformed cells were grown at 37°C to an OD_600_ ∼0.6 and induced with 0.2 mM IPTG overnight at 18°C. Cells were harvested by centrifugation (5000 × *g* for 10 min), and bacterial pellets were resuspended in lysis buffer and flash-frozen in liquid nitrogen. Cells were lysed using a Branson digital sonifier and clarified by centrifugation (12,000 × g for 30 min).

For 6×His-tagged Mud^CC^, cells were lysed in N1 buffer (50 mM Tris pH8, 300 mM NaCl, 10 mM imidazole) and coupled to Ni-NTA resin (Thermo Fisher Scientific, catalog #88222) for 3 h at 4°C. Following extensive washing, protein was eluted with N2 buffer (50 mM Tris pH8, 300 mM NaCl, 300 mM imidazole). The 6×His tag was removed using TEV protease during overnight dialysis into N1 buffer. Cleaved product was reverse affinity purified by a second incubation with Ni-NTA resin and collection of the unbound fraction. Final purification was carried out using an S200-sephadex size exclusion column equilibrated in storage buffer (20 mM Tris pH8, 200 mM NaCl, 2 mM DTT).

For all MBP-tagged aldolase proteins, cells were lysed in Phosphate Buffered Saline (PBS), and lysate was then clarified by centrifugation (12,000 × g for 30 min) and immediately flash frozen in liquid nitrogen for storage at −80°C. Proteins were isolated by coupling to amylose resin followed by extensive washing prior to their direct use in pulldown assays (see below).

### Pulldown assays

MBP-fused aldolase proteins were absorbed to amylose agarose for 30 min at 4°C and washed three times in wash buffer (20 mM Tris, pH 8, 100 mM NaCl, 1 mM DTT, and 0.2% Triton-X100) to remove unbound protein. Subsequently, soluble Mud^CC^ prey protein was added at varying concentrations for 2 h at 4°C with constant rocking in wash buffer. Incubation for different times (e.g. 1 or 3 h at 4°C, or 1 h at room temperature) produced similar results, indicating that this experimental framework had established equilibrium binding conditions. Reactions were then washed four times in wash buffer, and resolved samples were analyzed by Coomassie blue staining of SDS-PAGE gels. All gels shown in figures are representative of at least 4 independent experiments.

All interactions were quantified using ImageJ software. Briefly, gel images were converted to greyscale and individual band intensities were measured using the boxed ‘Measure’ analysis tool. The size of measurement box was kept the same across all concentrations. To ensure consistent measurements of bound proteins across experimental conditions, the measurement box was set by the size of the MBP-fusion protein bait. Binding curves shown in figures plot measured band intensities (expressed as arbitrary units, ‘AU’) for bound Mud protein. Dissociation binding constants were calculated in GraphPad Prism using a one-site binding isotherm regression analysis. All plots and statistics were also performed in Prism.

### AlphaFold structural modeling

Protein structural modeling was performed using the AlphaFold server (www.alphafoldserver.com) utilizing the current AlphaFold 3 code (16). Primary protein sequences were input along with the desired copy number (e.g. ‘4’ for prediction and modeling of tetrameric assemblies). Confidence metrics were interpreted for model accuracy. Monomeric structures were assessed using predicted template modeling (pTM) scores; tetrameric structures were assessed using this along with the interface predicted template modeling (ipTM) scores, both of which assess the accuracy of the entire structure (17, 18). All models presented scored above the 0.6 threshold for likelihood, with most all scoring above the 0.8 threshold for high-quality predictions. Models were further interpreted and shown images were rendered in PyMOL (www.pymol.org).

### Sequence database analysis

Database searches were performed using BLAST (www.blast.ncbi.nlm.nih.gov). The following sequences were used as inputs for respective aldolase class searches: Class I aldolase enzyme (https://www.ncbi.nlm.nih.gov/protein/NP_001262985.1?report=genbank&log$=protalign&blast_rank=1&RID=M9M6K9CS013), Class II aldolase enzyme (https://www.ncbi.nlm.nih.gov/protein/EFE61766.1?report=genbank&log$=protalign&blast_rank=2&RID=M9MBFN75016), ‘full CT’ Class IIa aldolase enzyme (https://www.ncbi.nlm.nih.gov/protein/WP_085599007.1?report=genbank&log$=protalign&blast_rank=1&RID=M9MNMM3R016), ‘half CT’ Class IIa aldolase enzyme (https://www.ncbi.nlm.nih.gov/protein/WP_000440781.1?report=genbank&log$=protalign&blast_rank=1&RID=M9NB2A1X016), ‘no CT’ Class IIa aldolase enzyme (https://www.ncbi.nlm.nih.gov/protein/WP_010881293.1?report=genbank&log$=protalign&blast_rank=1&RID=M9NJDBMR016), and Class IIa/adducin aldolase domain (https://www.ncbi.nlm.nih.gov/protein/NP_001246421.1). Class I and II enzymes are highly divergent in sequence, easily allowing distinctions of the Class I subgroup (13). Class II enzymes were confirmed by AlphaFold modeling and structural alignment to that of the query sequence. For all sequences inferred as class IIa enzymes, identified proteins were not part of larger sequences typical of Adducin genes and were subsequently modeled using AlphaFold with a Zn^2+^ ion to demonstrate predicted binding at the conserved histidine triad conserved within the active site of bona fide class II aldolase enzyme (19). Class IIa/adducin domains were identified as those located at the N-terminal region of proteins also containing sequence C-terminal to the aldolase sequence that most likely constitutes additional Adducin-specific function (10). Phyla searched were selected as those representing the three kingdoms forming the tree of life along with additional ones at significant positions in animal evolution (20). All multiple sequence alignments were performed using ClustalW (https://www.genome.jp/tools-bin/clustalw), and images shown were constructed in SnapGene software (https://www.snapgene.com).

## RESULTS

### Structural modeling of Hts^ALDO^ domain reveals a distinct loop insertion in its CThelix

The recently determined cryo-EM structure of Adducin bound to a spectrin-actin complex found a hetero-tetrameric α/β-Adducin ALDO^DOM^ associated with the barbed actin end (PDBid: 8IAH; (11)). This tetrameric assembly, as well as folding of individual monomers, shares high homology with those of a subgroup of bacterial aldolase enzymes (e.g. PBDid: 3OCR; RMSD=0.735 for tetramer alignment; Figure 1). To examine how the ALDO^DOM^ of Hts compares, we constructed a tetrameric model using AlphaFold 3 (AF3; (16)), with an input sequence of Hts amino acids (residues 110-401) that align with those that constitute the complete aldolase fold in α-Adducin (11). AF3 produced a tetrameric Hts assembly with a high overall confidence (ipTM=0.87; pTM = 0.88), showing a majority of positions within the ‘Very high’ (pIDDT > 90) confidence range apart from an extended loop within the final, ‘C-terminal helix’ (CThelix) of each aldolase domain (Figure 1A). This model showed high homology to both the mammalian Adducin domain and bacterial enzyme tetramers (Figure 1B,C). We conclude that the Hts ALDO^DOM^ forms a tetrameric assembly with high structural homology to related aldolase fold proteins.

**Figure 1.**
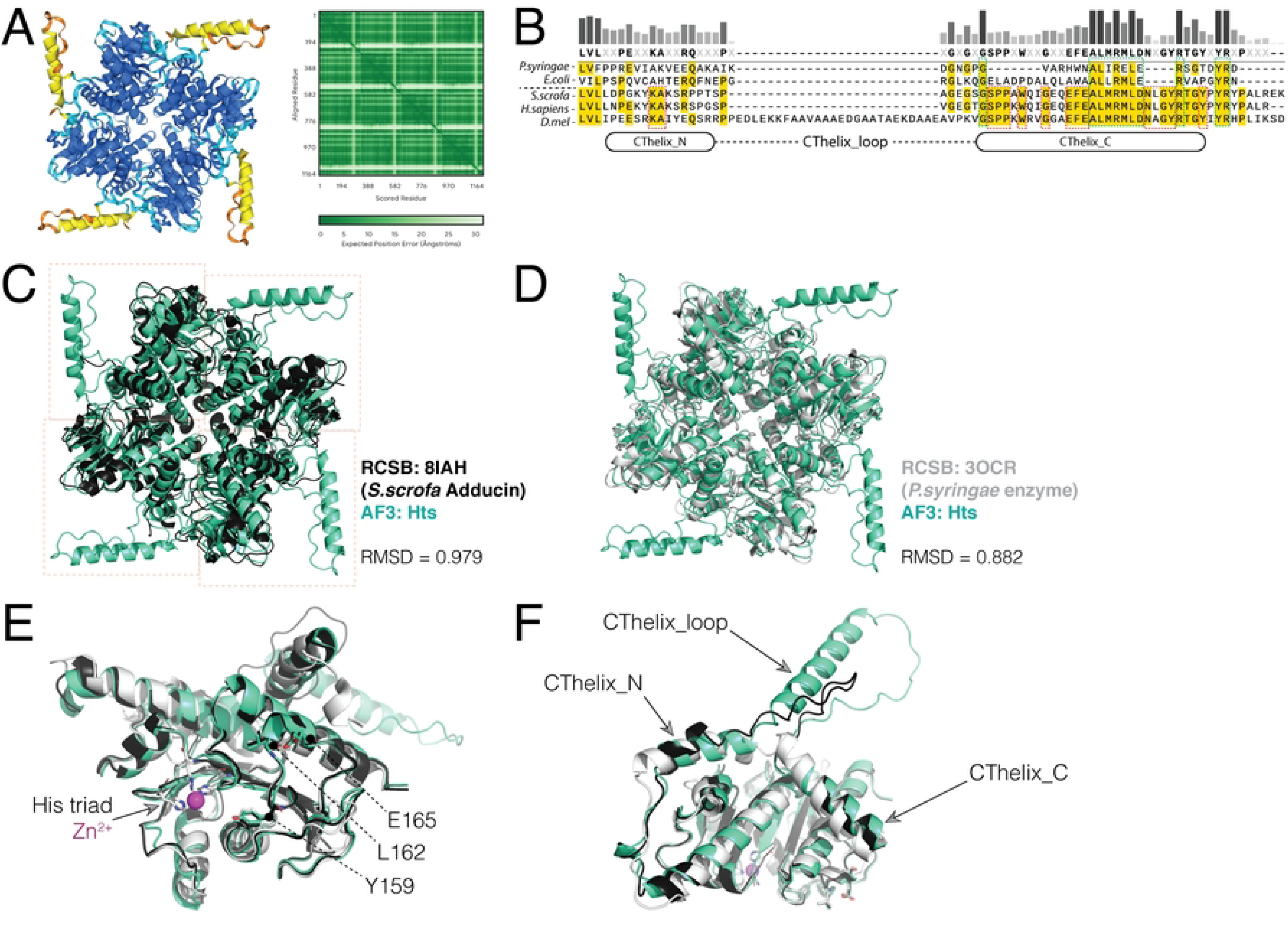
AlphaFold modeling indicates Hts contains an extended CThelix loop and forms a tetrameric assembly. A. *Left*: AF3 modeling output results for the Hts tetramer color coded by per-atom confidence estimate (pLDDT) on 0-100 scale (*Blue* – Very high: pLDDT > 90, *Cyan* – Confident: 90 > pLDDT > 70, *Yellow* – Low: 70 > pLDDT > 50, *Orange* – Very low: pLDDT < 50). The core aldolase fold in each of the four Hts monomers contains a majority of ‘Very high’ scoring positions, with a minor fraction of ‘Confident’. In contrast, the extended CThelix loop showed scores in the ‘Low’ and ‘Very low’ categories. *Right*: Predicted alignment error plot shows the expected positional error for each residue, also indicating a high confidence level (dark green) in model accuracy apart from the CThelix loop positions (light green). B. Multiple sequence alignment of CThelix sequences from bacterial aldolase enzymes (*Pseudomonas syringae* and *Escherichia coli*) and aldolase domains from pig (*Sus scrofa*, *S.scrofa*), human (*Homo sapiens*, *H.sapiens*), and fruit fly (*Drosophila melanogaster*, *D.mel*) demonstrates the extended insert sequence in fly Hts domain that occurs between the N- and C-terminal regions of the CThelix. Also indicated are positions conserved among enzymes and domains (*dashed green boxes*) and those only conserved among domains *(dashed red boxes*). C. Overlay of fly Hts (*bluegreen*) and pig Adducin (*black*) tetramers shows a high degree of similarities, with an RMSD = 0.979, calculated from a superimposition alignment in PyMOL. D. Overlay of fly Hts (*bluegreen*) and bacterial aldolase enzyme (*white*) tetramers shows a high degree of similarities, with an RMSD = 0.882. E. Overlay of Hts, Adducin, and aldolase enzyme monomers indicating conserved YLE motif positions (numbered according to fly Hts) that are positioned adjacent to the bacterial enzyme active site containing the histidine triad (‘His triad’) and catalytic Zn^2+^ (*magenta*). F. Overlay of Hts, Adducin, and aldolase enzyme monomers demonstrating the conserved structure of the N- and C-terminal segments of the CThelix (‘CThelix_N’ and ‘CThelix_C’, respectively) as well as the divergent loop insert segment (‘CThelix_loop’).

As mentioned, each Hts domain monomer has a large, conspicuous loop within its CThelix (Figure 1). Alignment of single monomers demonstrated that Adducin has a smaller loop insertion, and the bacterial enzyme has no significant loop but instead a short break in an otherwise contiguous CThelix (Figure 1B-D). AF3 modeling of the bacterial enzyme was able to accurately place the active site catalytic Zn^2+^ ion (Figure 1D). In contrast, neither Adducin nor Hts modeling showed such placement, consistent with divergence in the catalytic site and loss of enzymatic function in transition to aldolase domains (10). Overall, these results demonstrate the conserved nature of the core aldolase fold, highlight a marked difference in the CThelix, and substantiate the accuracy of AF3 modeling for both enzymes and domains.

### The CThelix loop regulates Mud^CC^ direct interaction with the Hts^ALDO^ domain

Having identified the CThelix as the most structurally divergent element in aldolase proteins, we next inspected the role for this Hts sequence in regulating Mud binding. To do so, we immobilized Maltose-binding protein (MBP)-fused Hts proteins on agarose resin and co-incubated with purified Mud^CC^ domain, which is responsible for direct Hts interaction (Figure 2A). The full-length Hts ALDO^DOM^ strongly bound Mud with low-micromolar affinity, similar to our previous studies (Figure 2B; (15)). Deletion of the large CThelix loop insertion resulted in a significant loss of Mud binding, as quantified by a reduction in the maximal binding (B_max_), with minimal effects on the affinity of this diminished interaction efficacy (Figure 2C). This result suggests that Mud binding could occur directly within this loop insertion. However, truncation of the entire CThelix, including the loop insertion, surprisingly restored complete Mud binding comparable to the full-length ALDO^DOM^ (Figure 2D), indicating that the binding site occurs within the minimal aldolase core fold. Taken together (Figure 2E), these data suggest a possible regulatory role for the CThelix in Mud binding, with the loop insertion playing an important but indirect part of the interaction.

**Figure 2.**
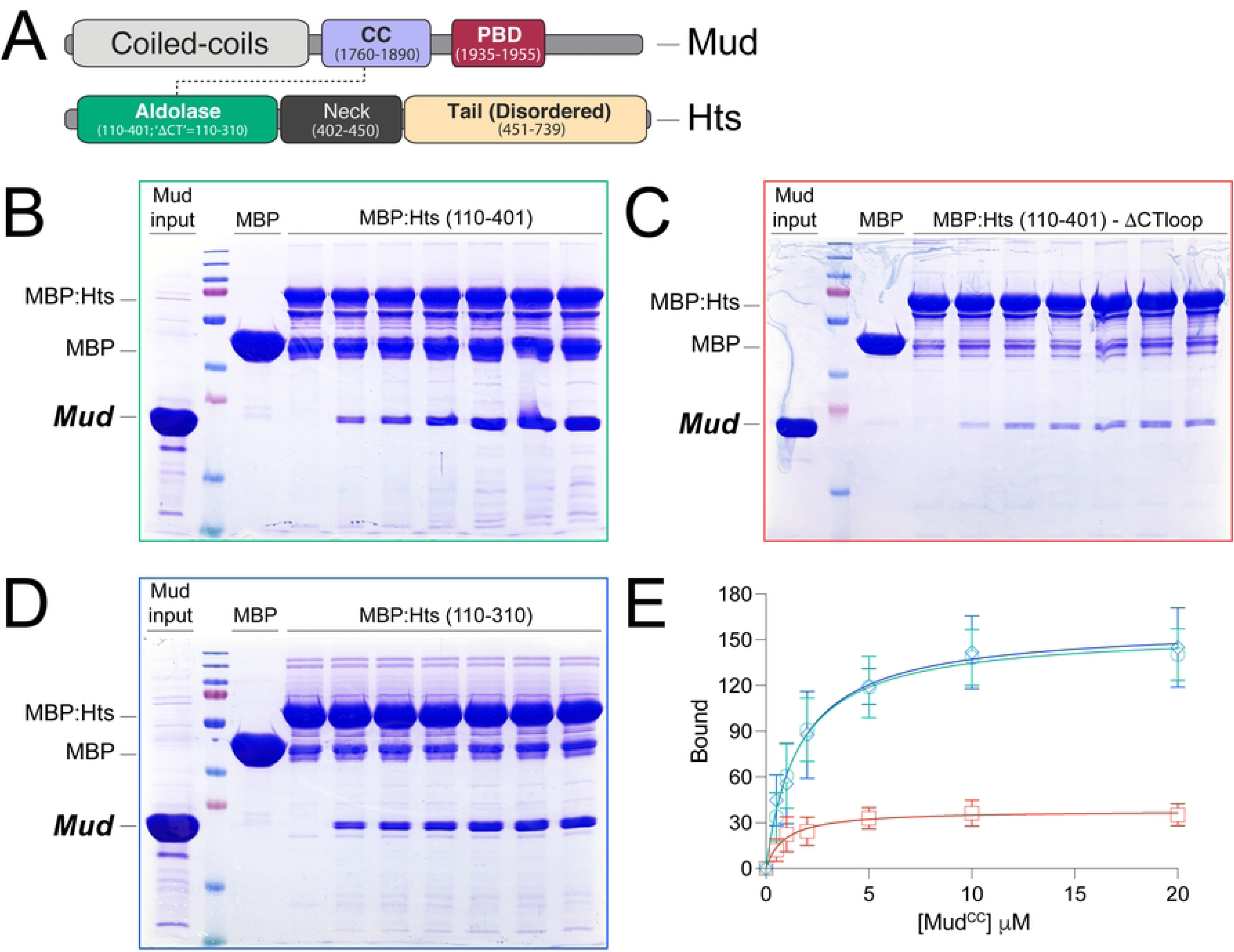
The Hts aldolase domain CThelix regulates its direct interaction with Mud^CC^. A. Domain diagrams for Mud and Hts proteins are shown. Mud (*top*) contains extended coiled coils at the N-terminus (*grey*), along with a shorter, more C-terminal positioned coiled coil (CC, *purple*) that is the focus on the current study. This Mud^CC^ domain is followed by the Pins-binding domain (PBD, *red*). Hts (*bottom*) contains the N-terminal Aldolase domain (*bluegreen*) that is also the focus of this study, followed by the Neck (*grey*) and disordered Tail (*light orange*) regions. B. Representative gel image depicting interaction between the full-length Hts aldolase domain and Mud^CC^. MBP-fused Hts [MBP:Hts (110-401)] or MBP alone control were coupled to amylose resin and incubated in the absence or presence of increasing concentrations of Mud^CC^. Reactions were resolved by SDS-PAGE and stained with Coomassie blue. Gel shown is representative of 4 independent experiments. C. Representative gel image depicting interaction between a CThelix loop deletion in the Hts aldolase domain and Mud^CC^. MBP-fused Hts [MBP:Hts (110-401)-ΔCTloop] or MBP alone control were coupled to amylose resin and incubated in the absence or presence of increasing concentrations of Mud^CC^. Reactions were resolved by SDS-PAGE and stained with Coomassie blue. Gel shown is representative of 4 independent experiments. D. Representative gel image depicting interaction between a CThelix truncation in Hts aldolase domain and Mud^CC^. MBP-fused Hts [MBP:Hts (110-310)] or MBP alone control were coupled to amylose resin and incubated in the absence or presence of increasing concentrations of Mud^CC^. Reactions were resolved by SDS-PAGE and stained with Coomassie blue. Gel shown is representative of 4 independent experiments. E. Saturation binding curves for all the above experiments demonstrate the effects of CTloop deletion and CThelix truncation mutants on Mud^CC^ binding to the Hts aldolase domain. Full-length Hts domain (*green*) shows a robust, dose-dependent binding. Deletion of the CTloop (*red*) significantly impairs Mud binding, whereas truncation of the entire CThelix (*blue*) restores strong binding indistinguishable from the full-length domain.

### Mutations at conserved surface residues on the Hts tetramer do not impair Mud binding

We next sought to identify a specific Mud binding site within the Hts aldolase domain. The ability of Mud to bind following removal of the Hts CThelix narrows the binding site to a minimal aldolase fold (residues 111-310), eliminating the small number of conserved contacts between the Adducin CThelix and actin as likely contributors. The CThelix loop insertion is notably disordered in the Adducin/Actin complex (11), consistent with it not being direly involved similar to our results above with Hts/Mud binding. Furthermore, a majority of Adducin contacts with Actin involve residues in extended sequences N- and C-terminal to the minimal aldolase core (11), with D339 (D310 in Hts) representing one exception. These finding together suggest that Mud binding occurs at a unique site within Hts. To examine this directly, we constructed a single alanine mutant at the conserved D310, along with three additional double mutants that were identified as conserved residues on surface-exposed regions of the Hts core tetramer (Q151A/E152A, Q177A/V181A, R205A/D207A; Figure 3A). Saturation binding experiments found that none of these mutants impaired Mud binding relative to a wild-type control Hts (Figure 3B-C). These data suggest that Mud interacts with a distinct site within the Hts ALDO^DOM^ that lies outside of conserved regions of the tetrameric complex surface.

**Figure 3.**
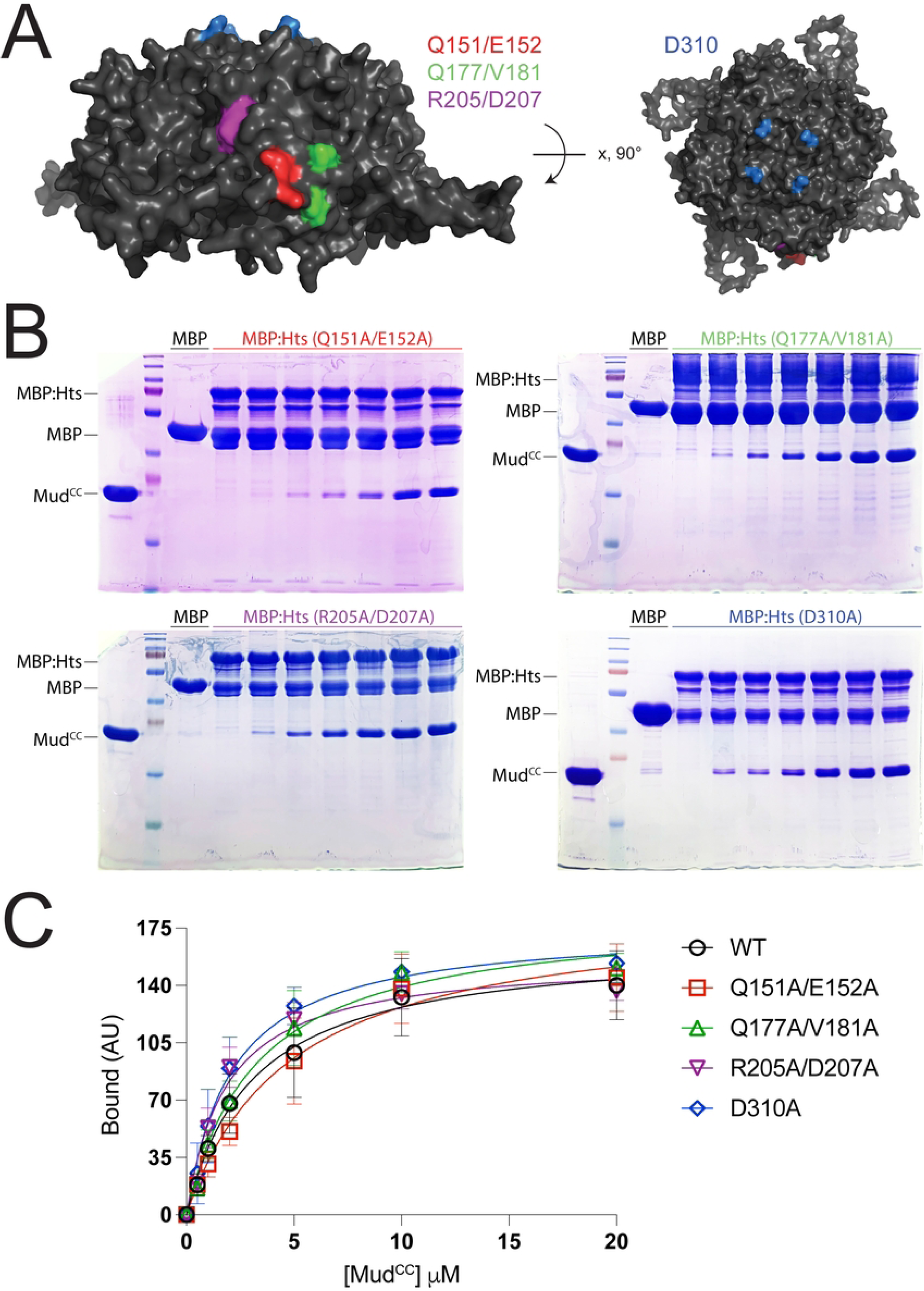
Conserved surface and actin-binding Hts residues do not contribute to Mud binding. A. *Left*: Lateral view of Hts tetramer highlighting three conserved surfaces residue pairs targeted for mutational analysis. *Right*: Rotation depicting the tetramer face with the conserved actin-contacting D310 residue also targeted for mutational analysis. B. Representative gel images depicting interaction between in the Hts aldolase domain mutants and Mud^CC^. Each indicated MBP-fused Hts mutant or MBP alone control were coupled to amylose resin and incubated in the absence or presence of increasing concentrations of Mud^CC^. Reactions were resolved by SDS-PAGE and stained with Coomassie blue. Gels shown are representative of 4 independent experiments. C. Saturation binding curves for Hts aldolase domain wild-type and mutants binding Mud^CC^. All mutants showed similar binding dynamics compared to control Hts.

### An ancient class II aldolase enzyme binds Mud upon CThelix truncation

We next explored Mud binding to a primitive aldolase enzyme from *Pseudomonas syringae*, whose CThelix lacks the loop insertion found in Hts and α-Adducin domains but nevertheless makes similar contacts to its adjacent protomer within an otherwise conserved tetrameric assembly (Figure 1D and 4A). This ALDO^ENZ^ lacked significant Mud binding, a result not unexpected considering its suspected role as a metabolic enzyme (Figure 4B-C). Remarkably, truncation of the CThelix from this ALDO^ENZ^ conferred robust Mud binding (Figure 4B-C), which was nearly equivalent to that seen with the ALDO^DOM^ of Hts, both ‘full length’ and ΔCThelix. AF3 modeling indicated that CThelix removal from the *Pseudomonas* enzyme does not prevent tetramer assembly (Figure 4A), consistent with the known tetramerization of aldolase enzymes naturally lacking a CThelix (see below). To corroborate this finding, we performed similar experiments with the homologous aldolase enzyme from *Escherichia coli*. We found this enzyme also lacked Mud binding in its full-length, wild-type form, but showed a robust interaction following truncation of its CThelix (Figure 4D).

**Figure 4.**
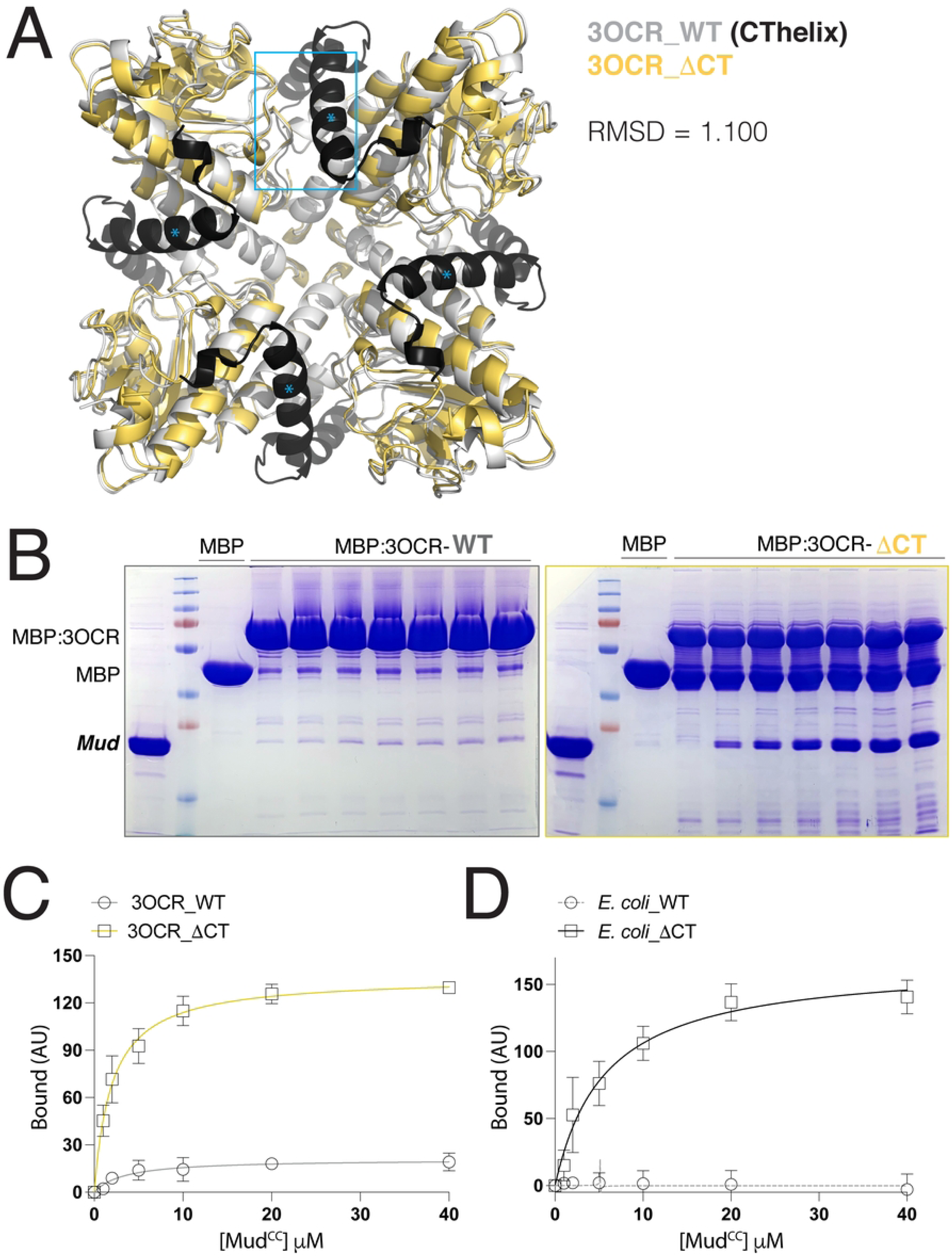
CThelix removal confers Mud binding function to a bacterial aldolase enzyme. A. Overlay of the full-length *P. syringae* aldolase enzyme (PDBid: 3OCR; *white* with CThelix colored *black*) tetramer with that of an AF3-generated model following truncation of the CThelix (ΔCThelix; *yellow*). The ΔCThelix enzyme retains a confident prediction for tetrameric assembly (pTM score = 0.7; ipTM score = 0.62) and aligns well to the full-length enzyme structure (RMSD = 1.1). Cyan box and asterisks depicts interaction of the CThelix (*) with loops in the aldolase core of its adjacent protomer. B. Representative gel images depicting Mud^CC^ interaction with full-length (*top*) and ΔCThelix (*bottom*) aldolase enzyme from *P. syringae*. Each indicated MBP-fused aldolase enzyme or MBP alone control were coupled to amylose resin and incubated in the absence or presence of increasing concentrations of Mud^CC^. Reactions were resolved by SDS-PAGE and stained with Coomassie blue. Gels shown are representative of 4 independent experiments. C. Saturation binding curves for conditions described above. Whereas the full-length aldolase enzyme shows only minimal Mud-binding capacity, truncation of the CThelix results in a robust Mud interaction.

We next constructed two chimeric aldolase proteins, swapping the CThelices between the Hts ALDO^DOM^ and the *Pseudomonas* ALDO^ENZ^, to further investigate the impact of this divergent structural element in Mud binding (Figure 5A). Compared to the wild-type enzyme that lacks significant Mud binding, chimeric fusion with the Hts CThelix resulted in a strong interaction similar to that seen following truncation of the enzyme CThelix (Figure 5B). In contrast, fusion of the Hts domain with the CThelix from the bacterial enzyme reduced Mud binding affinity compared to the wild-type domain (Figure 5C). This inhibitory effect of the enzyme CThelix on Hts function was weaker compared to the strong gain-of-function seen in the reciprocal chimera. Taken together, these results further highlight a key regulatory role for the CThelix in controlling the Mud binding activity within the aldolase fold.

**Figure 5.**
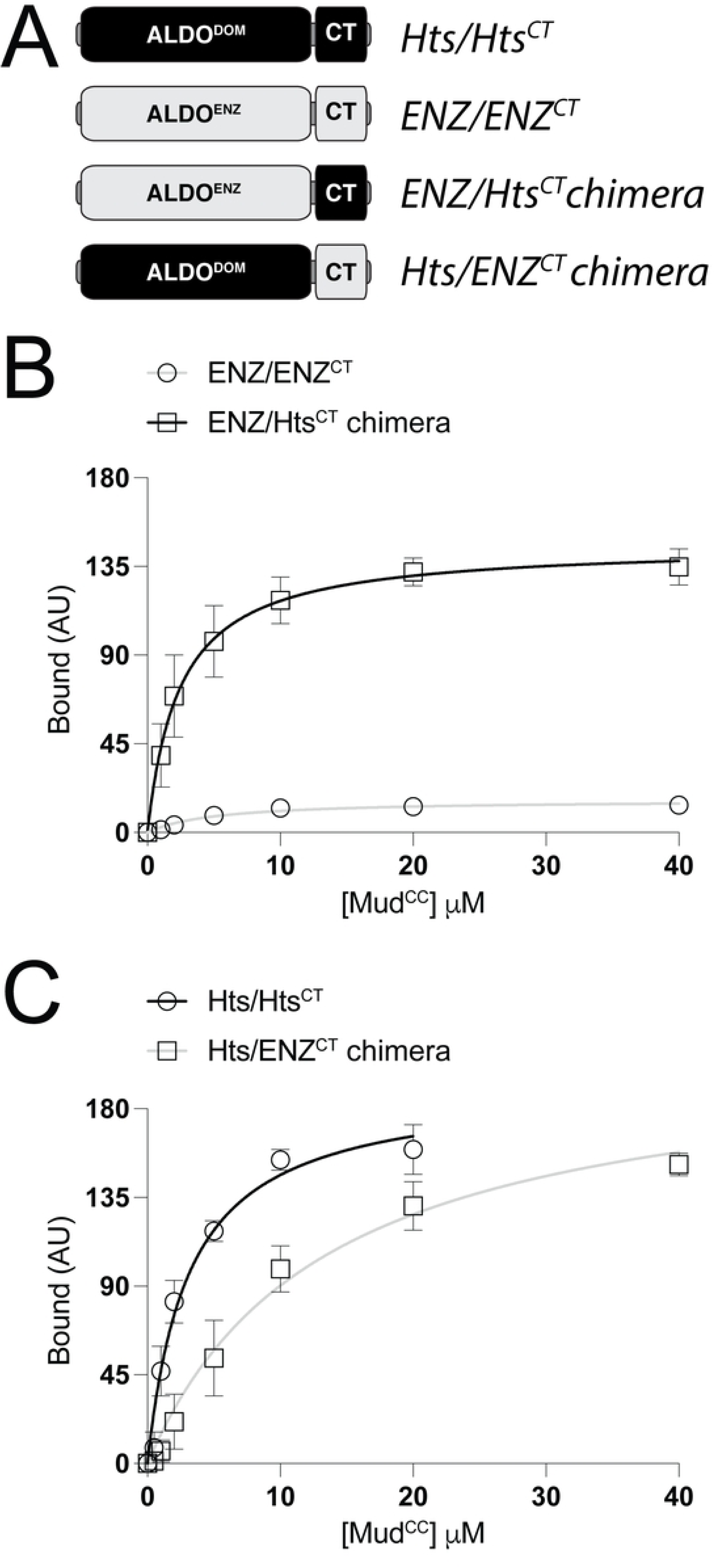
Chimeric enzyme/domain aldolase proteins suggest a key regulatory role for the CThelix in Mud binding function. A. Schematic representation of chimeric aldolase proteins shows strategy to swap CThelices (‘CT’) between the *Pseudomonas* aldolase enzyme (*light grey*; ALDO^ENZ^) and the Hts aldolase domain (*black*; ALDO^DOM^). B. Saturation binding curves for the full-length aldolase enzyme (ENZ/ENZ^CT^) and its chimeric version with the Hts CThelix (ENZ/Hts^CT^). Whereas the full-length aldolase enzyme shows only minimal Mud-binding capacity, the chimeric aldolase shows a robust Mud interaction. C. Saturation binding curves for the full-length Hts aldolase domain (Hts/Hts^CT^) and its chimeric version with the aldolase enzyme CThelix (Hts/ENZ^CT^). The chimeric aldolase shows a moderately impaired Mud binding compared to the full-length Hts domain.

### Evolutionary conservation at tetrameric interface identifies a putative Mud binding site within the core aldolase protein fold

Considering the gain-of-function Mud binding following truncation or chimerism of the aldolase enzyme CThelix, we next considered that the binding site may occur in a location repressed by this sequence in the full-length enzyme. To explore this idea, we constructed a multiple sequence alignment of Hts with several aldolase enzymes and Adducin domains to identify conserved residues near protomer interfaces that contact the CThelix in tetrameric structures (Figure 6). This analysis revealed several such positions within interconnecting loops of the core aldolase fold that directly contact the CThelix of the neighboring protomer (Figure 6A). Close inspection of this interface shows a particularly remarkable conservation within a triad of residues (F88/L91/E94 in the *Pseudomonas* enzyme and Y159/L162/E165 in the Hts domain; termed the ‘YLE motif’ herein) that contact a series of CThelix residues, some of which display similar conservation along with additional residues that constitute unique contacts in the aldolase domain (Figure 6C).

**Figure 6.**
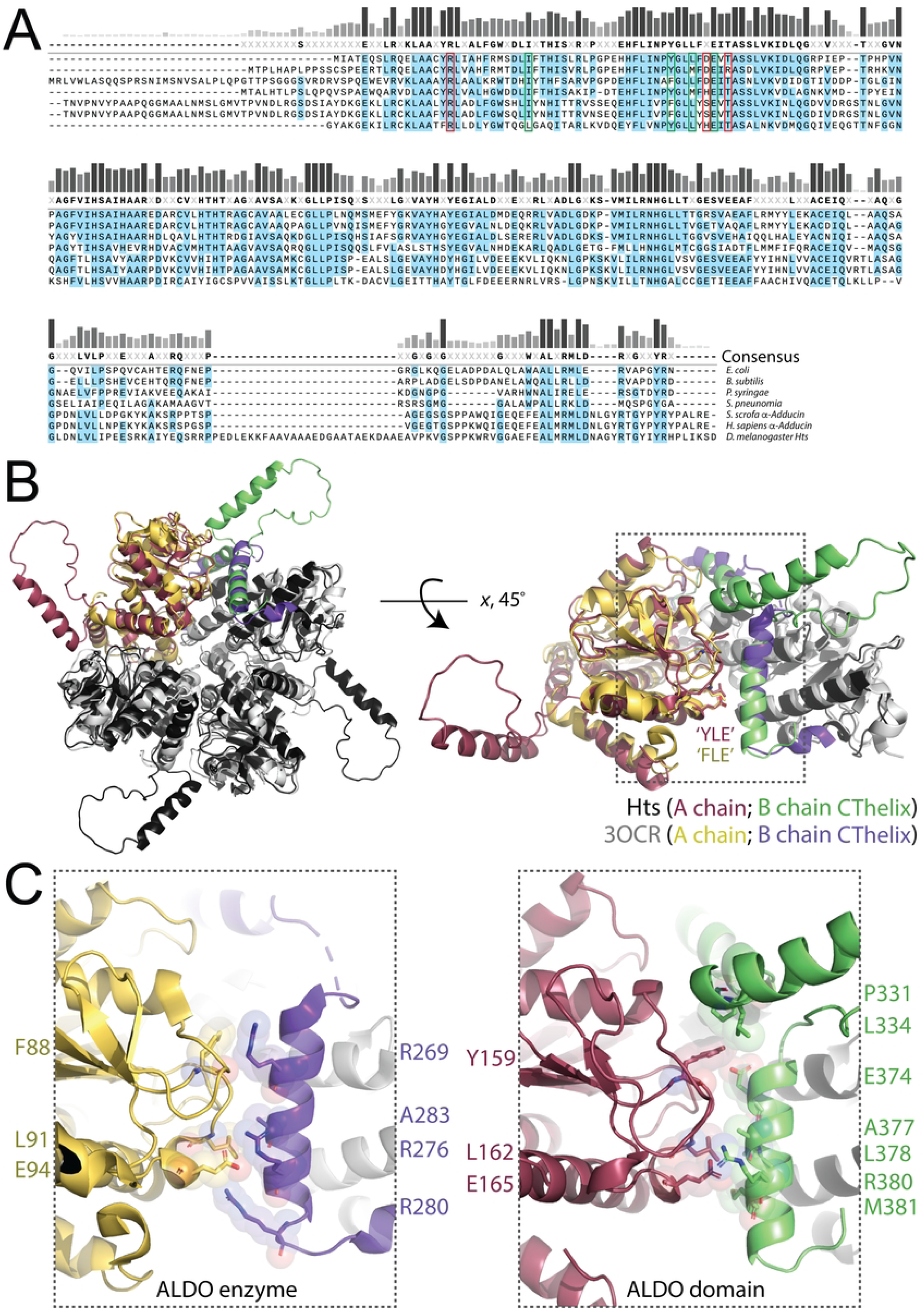
Hts shares a conserved YLE motif with primitive aldolase enzymes that contacts the adjacent protomer CThelix in the tetrameric assembly. A. Multiple sequence alignment among several bacterial aldolase enzymes and aldolase domains. *Green boxes* indicate conserved residues in the YLE motif, whereas *red boxes* indicate additional conserved residues that neighbor the YLE motif in the core aldolase structure. These residues were tested for contributions to Mud binding (see Figure 7). B. *Left*: face view of overlay of tetrameric assemblies for Hts aldolase domain (*black*; chain A is *red*, chain B CThelix is *green*) and the *Pseudomonas* aldolase enzyme (*white*; chain A is *yellow*, chain B CThelix is *purple*). *Right*: lateral view of contacts between chain A and B protomers with the conserved, CThelix-contacting YLE motif residues shown as sticks. C. Zoomed views of YLE-CThelix contacts in the aldolase enzyme (*left*; ALDO enzyme) and aldolase domain (*right*; ALDO domain) demonstrate the conserved nature of these interactions. Specific contacting residues are indicated.

To determine the role of these conserved positions in the Hts ALDO^DOM^, we constructed a series of alanine mutants and measured their Mud binding capacity relative to wild-type. Triple mutation of the YLE motif resulted in a complete loss of Mud binding when examined within the context of either the full-length domain or the minimal core (e.g. ΔCThelix; Figure 7A). A single L139A also resulted in a significant impairment of binding (Figure 7A). In contrast, double mutation of H164/T167 bound Mud similar to wild-type, as did a single R128A mutant (Figure 7B). These data highlight a critical role of the YLE motif in Mud binding and discount a role for the adjacent H164/T167 and R128 residues. To further examine the importance of the YLE motif, we tested the effects of mutations to the equivalent FLE motif in the *Pseudomonas* aldolase enzyme. When examined within the context of the gain-of-function ΔCThelix enzyme, this mutation resulted in significantly impaired Mud interaction, similar to the YLE mutation in Hts (Figure 7C). Overall, these data implicate an important role for the conserved YLE motif in Mud binding (Figure 7D).

**Figure 7.**
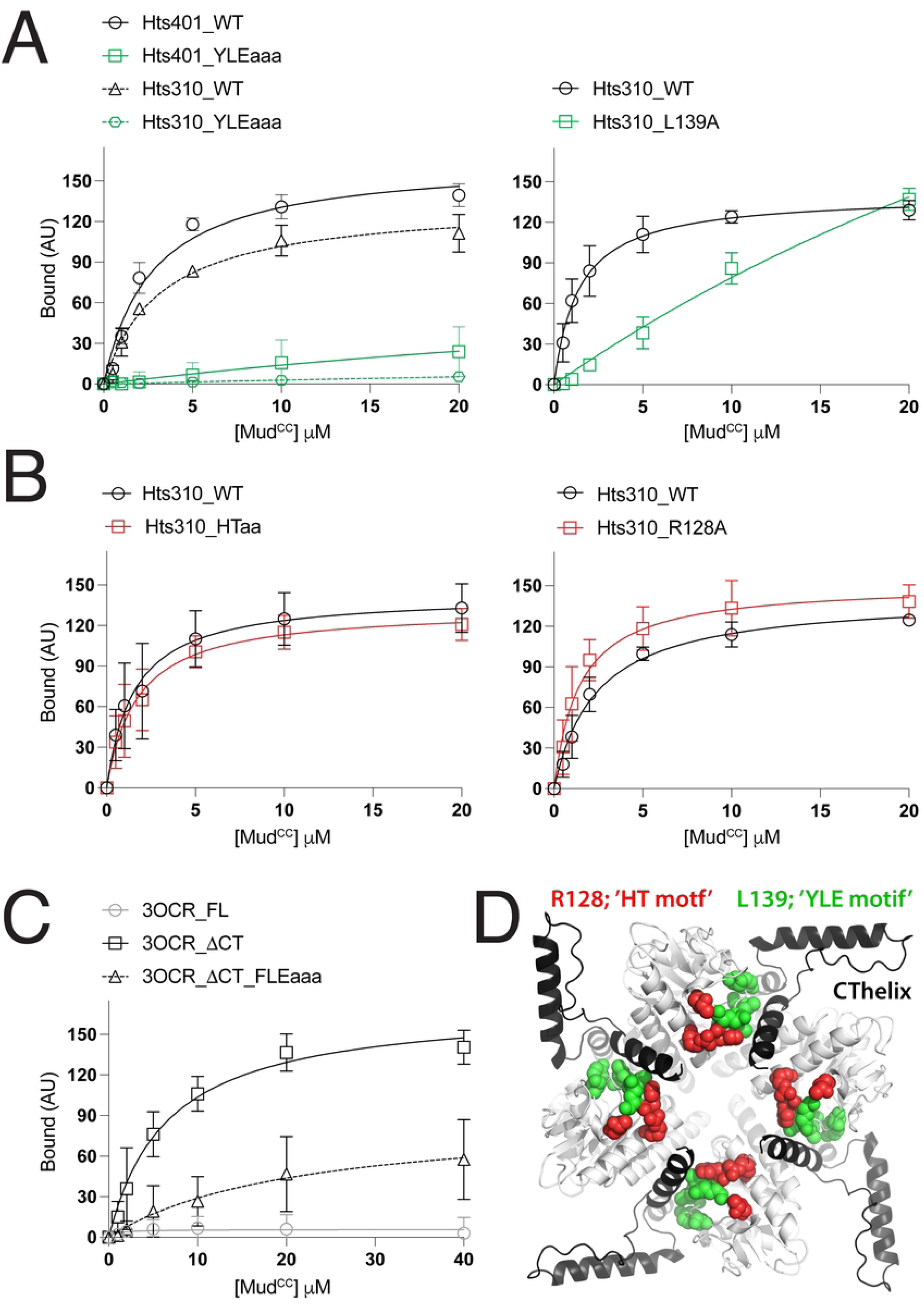
Mutation of the conserved YLE motif impairs Mud binding. A. *Left*: YLE residues were mutated as a triple-alanine mutant (YLEaaa) in either the full-length Hts aldolase domain (Hts401) or the core fold lacking the CThelix (Hts310) and tested for Mud binding compared to wild-type (WT). In both cases, the YLEaaa mutant significantly impaired Mud binding. *Right*: a single alanine mutation was made to the conserved L139 within the core aldolase domain (Hts310_L139A), which also negatively impacted Mud binding dynamics. B. *Left*: A double alanine mutation at two conserved positions adjacent to the YLE motif was made in the core domain (Hts310_HTaa). This mutant retained full Mud binding function. *Right*: Similarly, a single R128A mutation did not impair Mud binding. C. Triple-alanine mutation of the FLE motif residues in the *Pseudomonas* aldolase enzyme (3OCR) significantly reduced the gain-of-function Mud binding seen following truncation of the enzyme CThelix. The full-length (FL) enzyme again did not bind Mud. D. AF3-generated tetrameric assembly of the full-length Hts aldolase domain (*white* with CThelices colored *black*). YLE motif and L139 residues, those whose mutation impaired Mud binding, are shown in *green*, whereas the adjacent residues that were not required for Mud binding are shown in *red*.

### Aldolase enzyme subgroups with varying CThelix length display differential Mud binding capacity

Having identified a putative Mud interaction site conserved between the Hts aldolase domain and *Pseudomonas* enzyme, we next widened our analysis to additional aldolase enzymes. Searching structural and sequence databases, we noted three subgroups within the Adducin-like class II aldolase enzymes (those with structural homology to the aldolase domain, referred to as ‘class IIa’ herein), which could be distinguished by the length of their CThelix (Figure 8A-B). These enzymes mostly belong to a group of tetrameric L-fuculose-1-phosphate aldolases found in many pathogenic bacteria (21). The first subgroup completely lacks a CThelix (termed ‘no CT’ herein); nevertheless, they share a core structure and tetrameric assembly with those aldolases already discussed (e.g. PBDid: 2IRP; Figure 8B). Ostensibly, the 2IRP enzyme mimics the *Pseudomonas* enzyme following truncation of its CThelix. In contrast to this enzyme, however, we found that the ‘no CT’ enzyme was incapable of binding Mud (Figure 8C). Primary sequence comparison revealed that the ‘no CT’ enzyme is highly divergent in the core loop sequences, including the YLE motif (Figure 8A). Additional ‘no CT’ aldolase structures also demonstrate that, unlike ‘full CT’ enzymes, their divergent residues corresponding to the YLE motif do not make contacts with neighboring protomers in what are otherwise similar tetrameric assemblies (22). Mutation of these corresponding residues to YLE in the 2IRP enzyme did not lead to a gain of Mud binding function (Figure 8C), indicating this motif is necessary for Mud binding in relevant aldolase proteins but not sufficient for those naturally lacking this activity.

**Figure 8.**
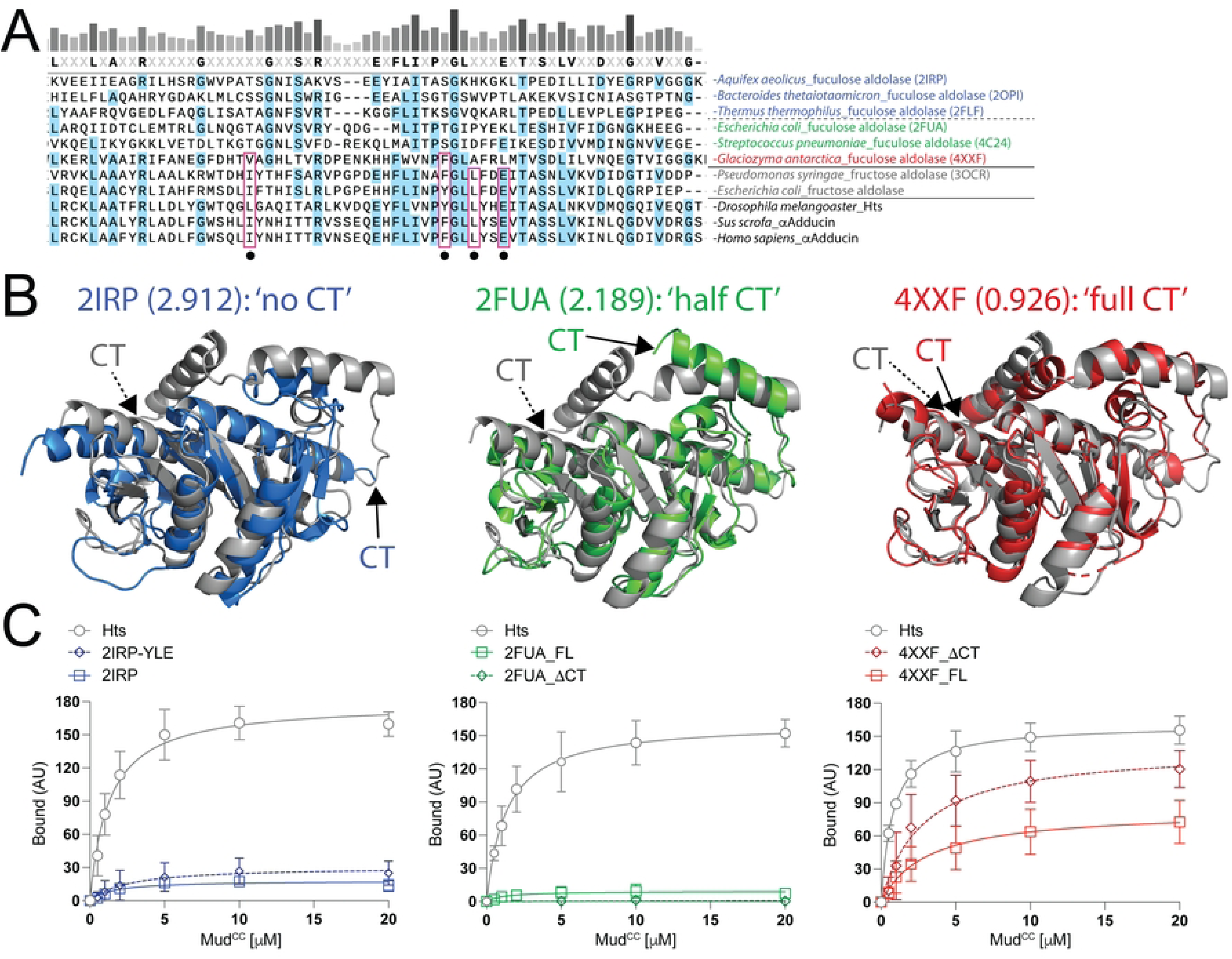
Divergent fuculose aldolase enzymes do not bind Mud despite their intrinsic lack of a complete CThelix. A. Multiple sequence alignment among several bacterial aldolase enzymes (with ‘aldolase’ label and respective PDBid codes where applicable) and aldolase domains (*black* text). Enzyme sequences shown include fuculose aldolases lacking a CThelix (*blue* text), those with a half CThelix (*green* text), and that with a full CThelix (*red* text), along with fructose aldolases with full CThelix sequences (*grey* text). Dots indicate positions corresponding to the YLE motif and L139 in the Hts domain, with those species showing conservation in these positions boxed in *red*. Note that these residues are divergent in fuculose enzymes, particularly those lacking a full CThelix. B. Images depicting structural superimpositions between the *Pseudomonas* fructose aldolase enzyme (3OCR; *grey* in each case) with indicated fuculose aldolases (colors corresponding to that described in panel A). RMSD values are shown for each. Fuculose aldolase enzyme monomers share structural homology with the fructose aldolase despite primary sequence divergence. C. Saturation binding curves for indicated fuculose aldolase enzyme interaction with Mud compared to the Hts aldolase domain. *Left*: the ‘no CT’ enzyme does not bind Mud despite its lack of a regulatory CThelix. Mutation of its divergent amino acids to YLE does not confer Mud binding. *Middle*: the ‘half CT’ enzyme also does not bind Mud, and truncation of its partial CThelix (ΔCT) does not confer binding. *Right*: the ‘full CT’ enzyme shows modest Mud binding relative to Hts and other fuculose enzymes. This binding is moderately improved upon truncation of the CThelix (ΔCT).

The second enzyme subgroup identified contains a short CThelix sequence (termed ‘half CT’ herein), which is structurally analogous to the sequence found prior to the loop insertion in Hts (Figure 8B). Structural studies have determined a similar tetrameric assembly in this enzyme subgroup as well (e.g. PDBid: 2FUA; (23)). As with the ‘no CT’ enzyme, we found that the ‘half CT’ enzyme was completely devoid of Mud binding. Furthermore, truncation of the shortened CThelix did not permit Mud binding (Figure 8C). Notably, as with ‘no CT’ enzymes, the ‘half CT’ aldolases also lacked conservation of the YLE motif (Figure 8A).

Lastly, database searches identified a third subgroup of fuculose aldolase enzymes, distributed across diverse fungi, that contain a CThelix of similar length to that of *Pseudomonas* and Adducin (e.g. PDBid: 4XXF; termed ‘full CT’ herein; Figure 8B). This ‘full CT’ 4XXF fuculose aldolase showed a moderate Mud binding activity that was improved upon truncation of its CThelix, although both responses were weaker than that measured with Hts (Figure 8C). Examination of its primary sequence found a partially conserved YLE motif (Figure 8A). Overall, these analyses further substantiate the CThelix and YLE motif as key molecular determinants for Mud binding in the aldolase protein fold.

### Evolutionary distribution of aldolase genes suggests a potential Placozoan emergence of the Adducin family

Aldolases represent a large family of surprisingly diverse enzymes with divergent sequences, structures, and reaction mechanisms that nonetheless contribute similar metabolic functions through convergent evolution (13). The starkest of these contrasts distinguish the class I and II subfamilies. Class I aldolases adopt a classic (and eponymous) TIM barrel fold and use a lysine-dependent Schiff base reaction mechanism (24, 25). Class II enzymes, in contrast, use a metal (most often Zn^2+^) catalyzed reaction scheme, with a conserved histidine triad functioning to coordinate this catalytic ion (19, 26). Class II enzymes can be further categorized structurally, with one subset also adopting a TIM barrel fold and a second showing high homology to the aldolase domain of Adducin proteins, along with other subgroups of sugar lyases (e.g. ‘class IIa’ enzymes; (27–29)). As discussed above, class IIa enzymes have a similar core structural fold but diverge in the length of their CThelices.

Considering these sequence and structural distinctions across the aldolase enzyme family, as well as the dramatic functional divergence between class IIa enzymes and the Adducin aldolase domain, we next extended our search of the protein sequence database (https://blast.ncbi.nlm.nih.gov/Blast.cgi) to identify expression patterns of these unique aldolase proteins across major phyla in an effort to further explore a potential pathway of functional evolution within this protein fold (Figure 9). We searched selected phyla for class I enzyme, class II enzyme, class IIa enzyme (‘no CT’, ‘half CT’, and ‘full CT’ versions), and Adducin-like aldolase domain sequences as depicted in Figure 9A. We found class I enzyme sequences present across all phyla apart from the *Choanoflagellates* (Figure 9B). Our searches also indicate that class I enzymes represents the lone subtype found across higher eukaryotes (with the exception of a class II sequence also identified in the protostome *Lophotrochozoa*), consistent with previous studies finding a loss of class II genes in these phyla (30–32). Class II enzymes were found in bacteria and most of lower eukaryotic phyla, typically together with class I sequences (Figure 9B). Class IIa enzymes were identified among similar phyla, albeit slightly more restricted than the class II sequences (Figure 9B). Searches for the aldolase domain found a restricted pattern of expression to metazoa, with the exception of *Porifera* in which no such sequence could be identified (Figure 9B). Rather, according to our searches, the aldolase domain appears to have emerged in *Placozoa*, which were also found to be the last phylum with apparent class IIa enzyme sequences, with both a ‘no CT’ and ‘full CT’ sequence identified (Figure 9B). Searches of several *Choanoflagellate* species, which have been shown to be a critical origin for other multi-domain scaffold proteins similar to Adducin (4, 33–36), revealed neither class IIa enzymes nor aldolase domain-containing Adducin sequences (Figure 9B). Overall, our sequence analysis supports existing theories of the aldolase enzyme family evolution and highlight a potential important role for *Placozoa*, a phylum of basal animals, in the class IIa and Adducin gene subgroups.

**Figure 9.**
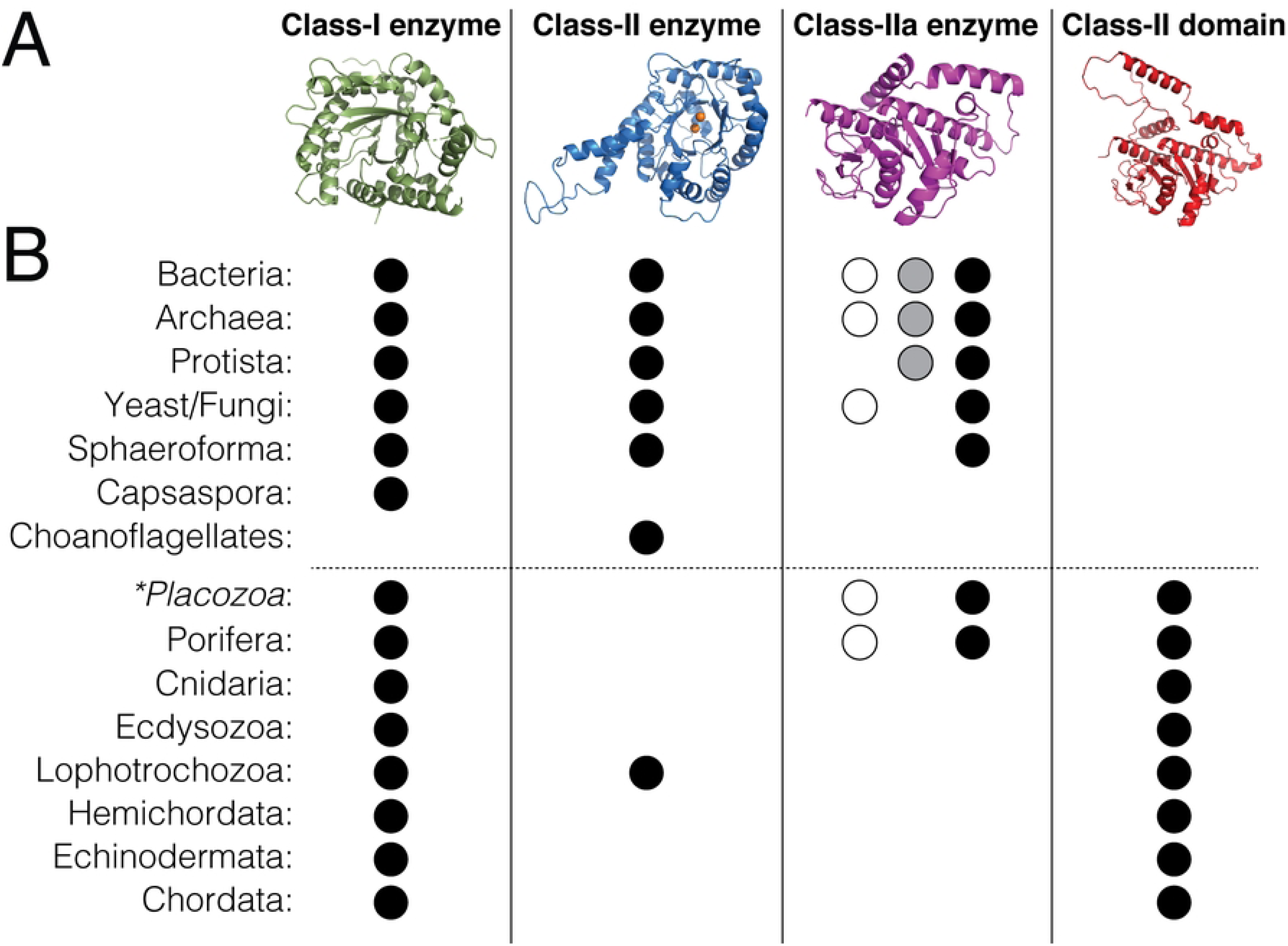
Sequence database analysis suggest a *Placozoan* origin for aldolase domain proteins. A. Structural representations of each subgroup of aldolase proteins are shown. B. Dots represent the positive identification of an annotated gene for the indicated aldolase protein in BLAST search analysis. For class IIa enzymes, the open dots represent ‘no CT’, grey dots represent ‘half CT’, and black dots represent ‘full CT’ sequences. Higher eukaryotes generally lack class II enzymes but gain expression of aldolase domain proteins (demarked by dashed horizontal line), whereas prokaryotes and lower eukaryotes lack domains and generally show a more diverse enzyme expression profile.

### A Class IIa aldolase enzyme from Trichoplax functions as a Mud binding protein

To empirically evaluate the *Placozoan* class IIa enzyme and domain genes identified, we began by modeling tetrameric structures of those from a prototype member, *Trichoplax adhaerens* (37). As shown in Figure 10A, AF3 predicted assemblies similar to those found for bacterial enzyme and Hts domain aldolase tetramers. The lower ipTM and pTM confidence scores for the domain assembly likely result from ‘Low’ confidence per-residue plDDT scores found in each protomer CThelix (not depicted). Notably, AF3 modeled Zn^2+^ ions within the conserved histidine triad retained in the enzyme sequence and structure, supporting the notion that these retain aldolase activity (Figure 10A-B). Similar metal placement was not predicted in the aldolase domain, which has diverged significantly within this catalytic triad (Figure 10B).

**Figure 10.**
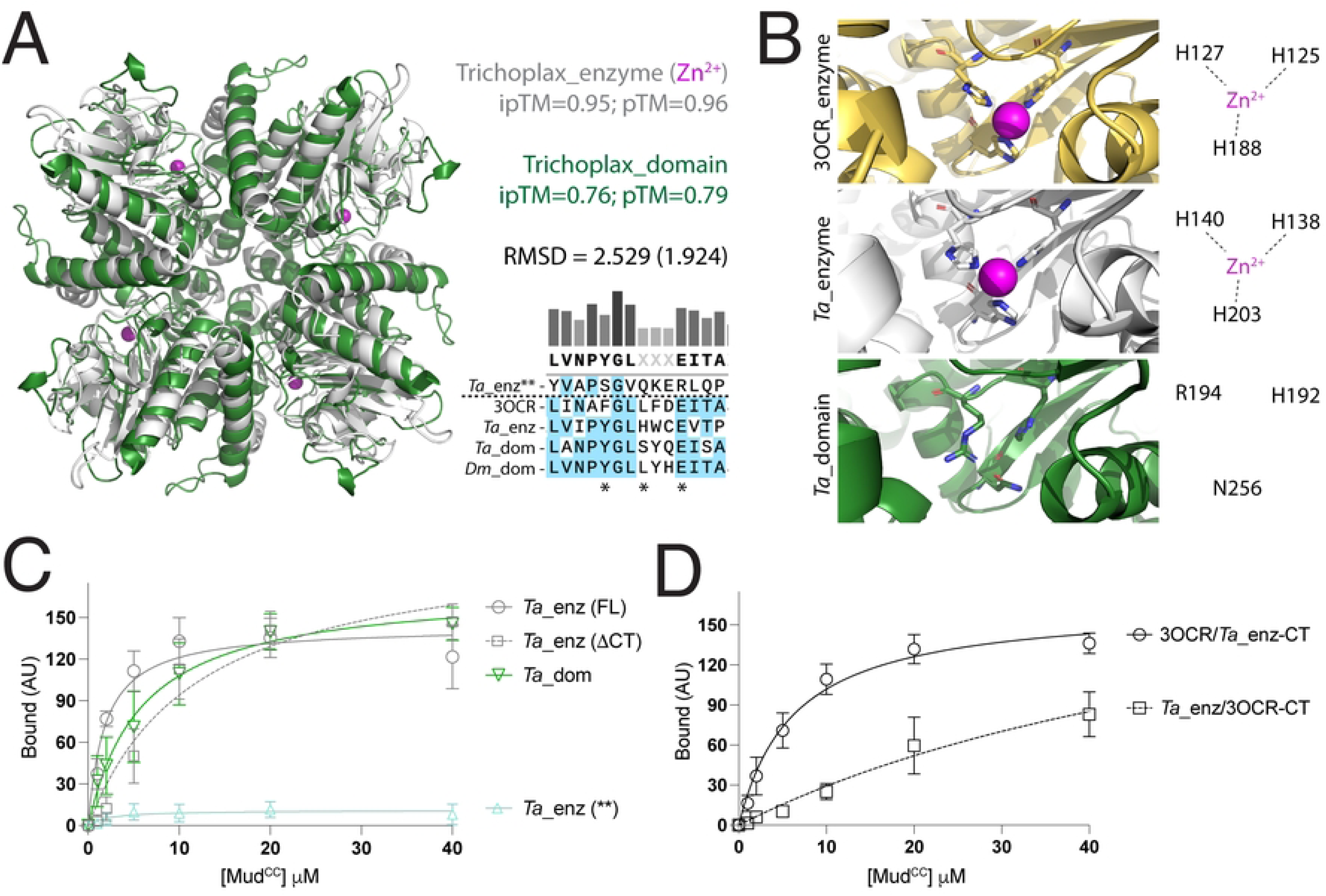
The *Placozoan* aldolase enzyme has full Mud binding activity and a CThelix with aldolase domain-like regulatory function. A. Overlay of AF3-generated tetramer models for the *T. adhearens* (*Ta*) aldolase enzyme (*white*) and Hts-like domain (*green*) shows similarities to other aldolase proteins. AF3 accurately models Zn^2+^ ions (*magenta*) into the conserved histidine triad within the enzyme catalytic site. Multiple sequence alignment shows conservation in the YLE motif (*) in the *Ta* enzyme and domain compared to fly Hts and *Pseudomonas* enzyme (3OCR). Sequence for a ‘no CT’ fuculose enzyme was also identified in *Ta* (enz**), which shows high sequence divergence in the YLE positions. B. Zoom views of the *Ta* aldolase enzyme (*top*), *Pseudomonas* aldolase enzyme (3OCR; *middle*), and *Ta* aldolase domain (*bottom*) shows the AF3-modeled Zn^2+^ in the *Ta* enzyme coordinated by the histidine triad nearly identical to the bacterial enzyme. In contrast, the *Ta* domain has diverged in two of these histidine residues and does not contain a catalytic metal. This is consistent with a retention of catalytic activity in the *Ta* enzyme. C. Saturation binding curves for the *Ta* aldolase proteins. Notably, the full-length (FL) enzyme containing a full CThelix binds Mud equally to the aldolase domain. Truncation of the enzyme CThelix does not improve or impair binding, similar to Hts (see Figure 2). In contrast, the divergent ‘no CT’ enzyme (**) does not bind Mud. D. Saturation binding curves for chimeric *Trichoplax* and *Pseudomonas* aldolase enzymes are shown. Fusion of the *Trichoplax* CThelix onto the core *Pseudomonas* enzyme (3OCR/*Ta*_enz-CT) shows robust Mud binding compared to the lack of interaction in the full-length bacterial enzyme (see Figure 4). In contrast, fusion of the *Pseudomonas* CThelix onto the core *Trichoplax* enzyme shows a moderate suppression of binding compared to the strong Mud binding seen in full-length *Ta* enzyme (compare to panel C).

We next examined the ability of the *Trichoplax* aldolase proteins to directly bind Mud in pulldown experiments. The *Trichoplax* ALDO^DOM^ bound Mud similarly to Hts, indicating that protein binding is a function present in Adducin proteins in the most basal metazoan (Figure 10C). Surprisingly, we found that the ‘full CT’ ALDO^ENZ^ from *Trichoplax* bound Mud to an equal extent, and truncation of its CThelix did not improve or impair binding (Figure 10C). Notably, this enzyme, as well as the identified aldolase domain, both contain a conserved YLE motif (Figure 10A). We also identified a ‘no CT’ enzyme in *Trichoplax*, which showed a divergent sequence at the YLE positions and was completely devoid of Mud binding function (Figure 10A,C).

Having found that the *Trichoplax* ‘full CT’ enzyme binds Mud at a level similar to the Hts aldolase domain, we next constructed enzyme chimeras with the *Pseudomonas* aldolase, which is devoid of Mud binding (Figure 4). Specifically, we swapped the CThelix between these enzymes and examined Mud binding ability. Chimeric *Pseudomonas* aldolase that has its CThelix replaced with that from *Trichoplax* was found to have a gain-of-function, showing Mud binding similar to that of the full-length *Trichoplax* enzyme (Figure 10D). In contrast, chimeric *Trichoplax* aldolase with its CThelix replaced with that from *Pseudomonas* was found to have impaired Mud binding relative to the wild-type enzyme (Figure 10D). These data further highlight a regulatory role for the CThelix in protein binding function of the aldolase fold.

### A single arginine-to-glycine change in an aldolase enzyme confers Mud binding

Truncation and chimerism of the CThelix, both of which lead to Mud binding in a primitive aldolase enzyme (Figures 4 and 5), represent relatively nontrivial sequence alterations. As such, we sought to identify changes that would have a minimal impact on aldolase enzyme sequence but a similarly large impact on Mud binding function. We considered two criteria to identify possible amino acid substitutions: (1) those located at or near protomer interfaces within the aldolase tetramer, and (2) those with significant distinctions between enzymes and domains conserved across evolution. This analysis revealed one residue in particular, which shows a strong conservation for arginine (or lysine; R/K) across several bacterial aldolase enzymes but that is substituted for glycine (G) in Adducin aldolase domains (Figure 11A). Although glycine was also identified in a subset of aldolase enzymes, it was unanimously present across domain sequences analyzed. Substitution of a cationic arginine for glycine represents a significant chemical change, and the strong conservation of these distinct amino acids further suggests a potential functional consequence. In both aldolase enzyme and domain structures, this residue is positioned immediately after the first α-helix of the aldolase fold, which packs against the CThelix (Figure 11B-C). In the aldolase enzyme, the arginine side chain points towards a central cavity formed in the tetrameric assembly. This results in the guanidinium R-groups of all four protomer arginine residues in the tetramer facing one another and packing closely together to form a narrow central cavity (8.6Å diameter; Figure 11B). In contrast, the lack of a non-hydrogen side chain in the aldolase domain glycine results in a loss of R-group packing and the formation of a significantly wider central cavity (19Å diameter; Figure 11C).

**Figure 11.**
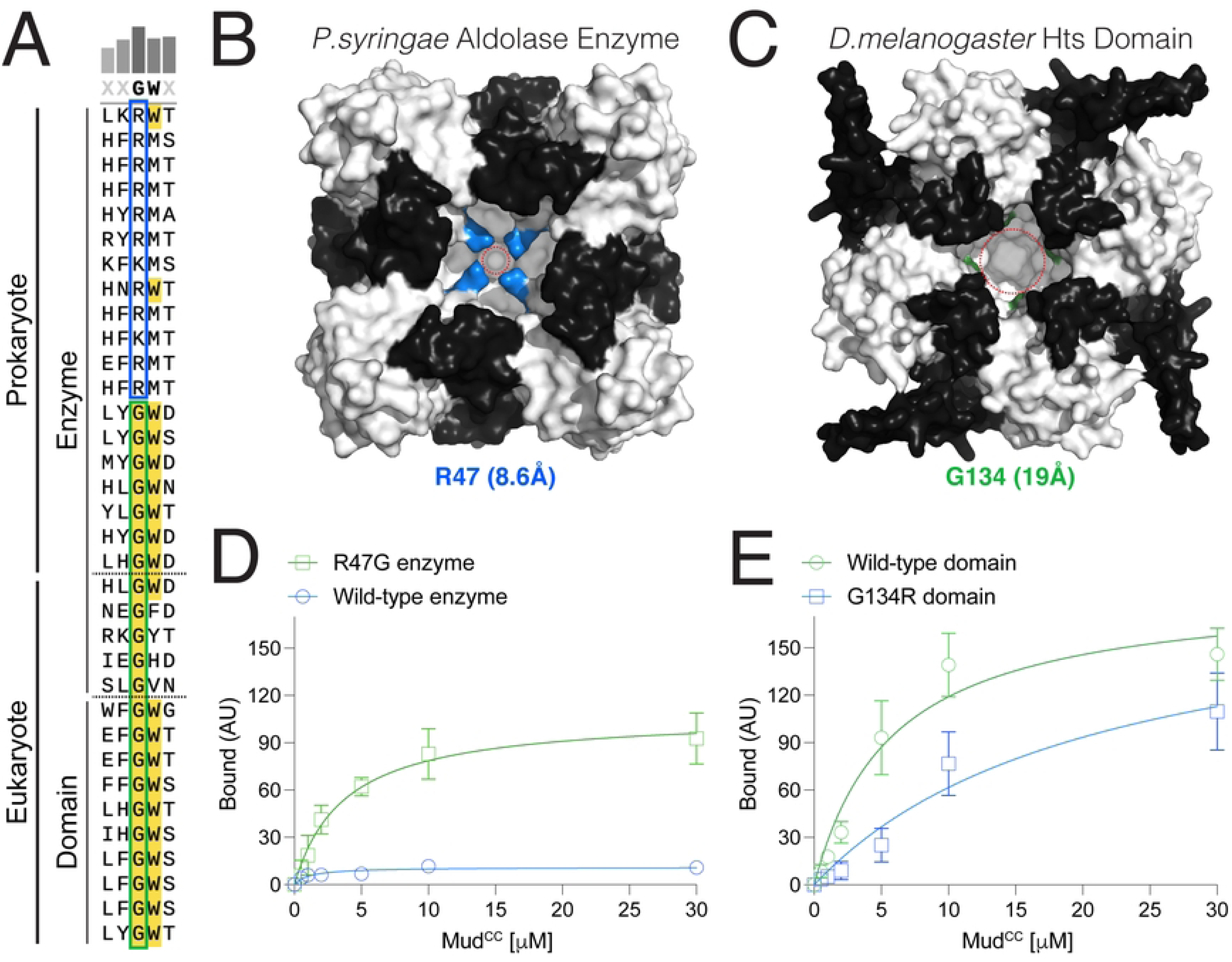
A single Arginine-to-Glycine substitution permits Mud binding in a bacterial aldolase enzyme. A. Multiple sequence alignment of aldolase enzyme and domain sequences demonstrates two distinct subgroups based on the highlighted position. A group of prokaryotic enzymes have a conserved arginine (R) or lysine (K) (*blue box*). Additional prokaryotic enzymes, along with primitive eukaryotic enzymes, have substituted this position with a glycine (G) (*green box*). Aldolase domains also have a well conserved glycine in this position. B. Tetrameric assembly of the *Pseudomonas syringae* aldolase enzyme (RCSBid: 3OCR) is shown in surface rendering with the main fold in *white* and CThelix in *black*. Arginine at position 47 (R47) in each protomer is colored *blue*, the side chains of which point toward a central core of the tetramer to create a cavity of 8.6Å diameter, denoted by the *dashed red* circle. C. AF3 model for the tetrameric assembly of *Drosophila melanogaster* Hts aldolase domain is shown in surface rendering with the main fold in *white* and CThelix in *black*. Glycine at position 134 (G134) in each protomer is colored *green*. The absence of a non-hydrogen side chain creates a cavity of 19Å diameter, denoted by the *dashed red* circle. D. Saturation binding curves for Wild-type (*blue*) and R47G (*green*) *Pseudomonas* aldolase enzymes are shown. The R47G mutant shows significant Mud binding compared to the wild-type enzyme. E. Saturation binding curves for Wild-type (*green*) and G134R (*blue*) *Drosophila* Hts aldolase domains are shown. The G134R mutant shows reduced Mud binding affinity compared to the wild-type domain.

To directly examine the role of this R→G change across aldolase proteins, we constructed an R47G mutation in the *Pseudomonas* enzyme and the reciprocal G134R substitution in the Hts domain and measured Mud binding compared to their respective wild-type controls. Mud binding was significantly enhanced in the single R47G aldolase enzyme substitution (Figure 11D). Binding to the R47G enzyme was not as strong as that seen following truncation of the enzyme CThelix, but nevertheless represents a significant gain-of-function. In contrast, Mud binding was reduced in the G134R substituted Hts aldolase domain, although this effect was more modest compared to the gain-of-function seen in the R47G enzyme (Figure 11E). These results agree with those from alterations in the CThelix, with domain-mimicking changes in the aldolase enzyme producing significantly stronger effects than enzyme-mimicking changes in the Hts aldolase domain. Overall, our results spotlight a single amino acid substitution that may represent a significant step in the evolution of aldolase protein function. It should be noted again that although many bacterial enzymes were found to contain an R/K, others were identified with a G at this position. The consequences of this substitution to the normal enzyme function, for example with respect to their catalytic activity, will require further investigation. Deviation from either R/K or G amino acids in this position was rarely seen across aldolase sequences, underscoring a unique functional role.

## DISCUSSION

Complex eukaryotic organisms are reliant on ubiquitous modular protein domains, many of which function as protein interaction platforms that combine to form multidomain scaffolds (38–41). Understanding how such domains emerged is critical to understanding their molecular function. Herein, we have explored the molecular determinants for protein binding function in the Hts/Adducin aldolase domain, which shares sequence and structural homology with a subgroup of extant class IIa aldolase enzymes involved in glycolytic metabolism (9, 10). Protein binding in the ALDO^DOM^ follows a dramatic functional transition from a well-defined primitive function (sugar lyase), substantiating it as an excellent case study for investigating neofunctionalization within a eukaryotic protein domain. Divergence at one or more catalytic residues conserved in the class IIa ALDO^ENZ^ explains the loss of enzymatic function in the Adducin family (10). Sequence changes that uniquely afford protein binding functionality are less clear, however.

Our results suggest that the CThelix plays a critical regulatory role in Mud binding within the aldolase fold. This conclusion is based on several notable observations: (1) the CThelix has undergone sequence and structural divergence between enzymes and domains, (2) the CThelix is dispensable for Mud binding to the Hts domain, but a seemingly domain-specific loop insertion is essential to binding when the CThelix is intact, (3) truncation of the CThelix affords Mud binding to an otherwise binding-deficient aldolase enzyme, and (4) chimeric fusion of CThelix sequences confers their respective regulatory properties on the reciprocal core aldolase proteins. Thus, Mud binding is a property intrinsic to certain aldolase enzymes that appears to be constrained by a noncompliant CThelix. It is also noteworthy that although the *Pseudomonas* enzyme lacked Mud binding (Figure 4), a yeast aldolase enzyme showed moderate binding that was further improved upon CThelix truncation (e.g. 4XXF protein; Figure 8B-C). Furthermore, the *Placozoan* aldolase enzyme demonstrated robust Mud binding indistinguishable from both *Placazoan* and *Drosophila* Hts domains (Figure 10C). These results are consistent with an evolutionary progression toward protein binding function. Also of note, removal of the large Hts CThelix loop impaired Mud binding, yet neither yeast nor *Placazoan* enzymes contain such an insert and have CThelices structurally more similar to the *Pseudomonas* aldolase (Figures 8B and 10A). Thus, this sequence element does not appear to be required for protein binding, but instead may provide additional regulatory control within the Hts/Adducin domain.

In contrast to this apparent regulatory role for the CThelix, our studies also identify the YLE motif as a putative binding site for Mud that lies within the aldolase core fold. This conclusion is based on several observations as well: (1) mutation of this YLE signature in Hts abolishes Mud binding independent of the CThelix, (2) mutation of the conserved FLE motif impairs the gain-of-function Mud binding seen following CThelix truncation in the *Pseudomonas* aldolase enzyme, and (3) ‘no CT’ aldolase enzymes across a range of phyla (from bacteria to *Placozoa*), which do not bind Mud despite their natural lack of a CThelix, are highly divergent in YLE sequence. It is also noteworthy that mutation of surface exposed residues on the aldolase tetramer did not affect Mud binding, consistent with an alternative binding site unique from that identified for the actin barbed end. The YLE motif is located at a tetrameric interface where it directly contacts the CThelix of an adjacent protomer (Figures 6 and 7D), which could explain the regulatory role of the CThelix as controlling exposure of the YLE motif. Such a model would require conformational plasticity in the tetramer as CThelix and Mud binding would be predicted to be mutually exclusive at the YLE motif. Evolutionary changes that impact substrate selection and catalysis by altering conformational dynamics are known for many classes of enzymes (42). Studies on aldolases have suggested conformational flexibility within the tetrameric assembly, although these have been restricted to class I enzymes (43, 44). To our knowledge, the conformational dynamics in class IIa enzymes and Hts/Adducin aldolase domains remain unexplored.

Aldolase expression patterns across major phyla reveal several notable conclusions: (1) the appearance of Adducin genes in higher eukaryotes coincides with singular expression of class I enzymes and loss of class II genes that are more often co-identified in lower eukaryotes and prokaryotes, (2) unlike their critical evolutionary position in the emergence of other multidomain scaffold genes, *Choanoflagellates* do not appear to represent a similar role for Adducin-related genes, and (3) instead, the basal metazoan phyla *Placozoa*, *Porifera*, and *Cnidaria*, which emerged 750-800mya (45), each have Hts/Adducin aldolase domain genes suggesting these non-bilaterian animals represent the first appearance of the Adducin family. In the cases of *Placozoa* and *Porifera*, class IIa enzyme genes were also identified, whereas *Cnidaria* appear to represent the most modern phyla to restrict aldolase enzymes to class I alongside their Adducin gene (Figure 9). The full capacity for Mud binding in the *Placozoan* enzyme is striking and is consistent with the functional evolution of the aldolase fold. Although its enzymatic activity has not been tested empirically, the histidine triad is conserved and predicted to coordinate a catalytic Zn^2+^ by AF3 modeling (Figure 10B). In contrast, the *Placozoan* aldolase domain has divergent sequence in the catalytic triad and does not bind Zn^2+^ in AF3 modeling. The domain is also a modular component of a larger Hts/Adducin gene, whereas the enzyme is a lone aldolase fold protein.

Overall, with respect to Mud protein binding function within class IIa aldolase enzymes, our work finds a lack of activity in bacteria, moderate activity in yeast, and full activity in *Placozoa* when compared to the native interaction with the *Drosophila* Hts domain. A complete perspective of this function within aldolase proteins across evolution will require further studies. We speculate that the protein-protein interaction function of the aldolase domain co-opted a tetrameric assembly surface following divergence within the regulatory CThelix that may have increased flexibility within the aldolase tetramer. As mentioned, this model necessitates conformational plasticity in the domain tetramer, as Adducin has been shown to exist primarily in this oligomeric state with little contribution of a monomeric form (46). Protein conformational dynamics often correlate with evolvability of new functions (47). The gain-of-function Mud binding seen in the R47G bacterial aldolase enzyme spotlights one potential route for such changes. In addition to widening a central cavity within the tetramer core due its simple hydrogen side chain (Figure 11B-C), this substitution might afford increased conformational dynamics due to the inherent flexibility of glycine positions in the protein backbone. Future studies investigating this possibility within aldolase proteins from additional phyla, along with direct examination of conformational flexibility within binding competent and incompetent aldolases, will be important.

Finally, it is relevant to compare the postulated mode of Mud binding to Hts against that of known Adducin interacting partners. The most well-known and characterized interaction with the Adducin ALDO^DOM^ is with the actin-spectrin complex. CryoEM analysis recently revealed that the tetrameric domain binds the actin barbed end using surface exposed residues, including within the CThelix, along with a more extensive network of residues located in extended sequences not part of the formal aldolase core (11). These extended regions lie outside the sequence boundaries examined in our studies with Mud and, thus, must not be involved. Similarly, truncation of the CThelix did not impair Mud binding, nor did alanine substitutions at conserved surface exposed residues, indicating these are also dispensable. Thus, the mode of ALDO^DOM^ interaction with Mud appears to be unique compared to F-actin. Adducin has also been shown to localize to spindle poles, independent of actin, through association with Myosin X via the aldolase domain (48). Notably, Adducin spindle association is regulated by CDK1 phosphorylation at a conserved serine within the CThelix, which is suggested to trigger conformational changes (48). Whether Myosin X interaction requires an intact CThelix or the conserved YLE motif is currently unknown. Exploring these open questions, along with identifying additional binding partners to the aldolase domain, will be important future steps in further understanding the function of this ancient protein fold.

## Notes

### Competing Interest Statement

The authors have declared no competing interest.

